# Challenging the Diffusion Barrier Paradigm: Biofilms Promote Flavin-mediated Electron Shuttling in *Shewanella oneidensis*

**DOI:** 10.64898/2025.12.04.692256

**Authors:** Yoshihide Tokunou, Yu Manabe, Nozomu Obana, Masanori Toyofuku, Nobuhiko Nomura

**Affiliations:** Department of Life and Environmental Sciences, University of Tsukuba, Tsukuba, Ibaraki 305-8577, Japan; Research Center for Macromolecules and Biomaterials, National Institute for Material Science, 1-2-1 Sengen, Tsukuba, Ibaraki, 305-0047, Japan; Degree Programs in Life and Earth Sciences, University of Tsukuba, 1-1-1 Tennodai, Ibaraki 305-8577, Japan; Microbiology Research Center for Sustainability, University of Tsukuba, Tsukuba, Ibaraki 305-8572, Japan; Tsukuba Institute for Advanced Research, University of Tsukuba, Tsukuba, Ibaraki 305-8572, Japan; Department of Medicine, Transborder Medical Research Center, University of Tsukuba, Tsukuba, Ibaraki 305-8577, Japan

## Abstract

Flavins are ubiquitous diffusible redox mediators that are crucial in enabling extracellular electron transfer (EET) in electroactive bacteria that often form biofilms in both natural and engineered environments. However, the behavior of flavin diffusion within biofilms remains poorly understood. In this study, we developed a colony-based electrochemical platform using interdigitated electrode arrays to quantify the diffusion coefficients of flavins within bacterial biofilms. We found that flavin diffusivity was about 11-fold enhanced in *Shewanella oneidensis* MR-1 biofilms than in the bulk solution. However, this enhancement was abolished in the gene deletion mutant that lacked the membrane-associated flavin-binding *c*-type cytochrome OmcA, which suggests that the flavin–OmcA interaction on cell surfaces facilitates flavin diffusion within the biofilm. Notably, the diffusion coefficients of other redox molecules, such as methylene blue and safranin, scarcely improved in biofilms, which validates the inference that flavin-specific interaction-accelerated diffusion occurs in biofilm environments. These findings uncover a phenomenon promoting long-range electron transfer, overturning the prevailing assumption that shuttling-based EET is hindered by slow molecular diffusion in biofilms. Our study highlights the functional importance of cell-surface cytochromes in overcoming the kinetic limitation of diffusion-based electron transfer, thereby shaping the bioenergetics in biofilms.

## Introduction

Extracellular electron transfer (EET) is a microbial process that delivers electrons generated from intracellular metabolism to extracellular electron acceptors such as metal oxides or electrodes ^1^. EET substantially influences element cycling on Earth and is of considerable interest in the fields of geobiology and geochemistry ^2^. In addition to their ecological relevance, EET-capable bacteria have been used in environmental and energy technology applications such as microbial fuel cells and microbial electrosynthesis ^3, 4^. Furthermore, the ubiquity of EET-capable bacteria across diverse ecosystems ^5, 6^ including the human gut ^7, 8^, oral cavity ^9^, and pathogenic environments ^10, 11^ has renewed interest in EET as a prevalent bacterial energetic system. Additionally, studying EET as an alternative form of energy conservation along with respiration and fermentation would provide new insights into microbial physiology and ecological strategies ^12, 13^.

In both natural and engineered environments, EET-capable bacteria often form multicellular aggregates known as biofilms. An estimated 40–80% of environmental bacteria exist as biofilms ^14, 15, 16^. Therefore, the molecular mechanisms underlying EET need to be elucidated within biofilms and not just in cells that are directly adsorbed onto electrodes. However, despite the significance of EET, our understanding of its operation in biofilm environments and the difference in its molecular mechanisms from that in electrode-adsorbed cells remains limited.

The involvement of flavin molecules as diffusible redox mediators constitutes a central component of EET in biofilms. In the Gram-negative model EET-capable bacterium *Shewanella oneidensis* MR-1, the secreted flavins shuttle electrons from the *c*-type cytochromes (e.g., OmcA and MtrC) located on the outer membrane to electrodes ^17, 18^. In Gram-positive bacteria such as *Listeria monocytogenes* and *Lactiplantibacillus plantarum*, flavins accept electrons from cell wall-anchored proteins and mediate electron transfer to extracellular acceptors ^12, 19^. EET occurs in the human gut bacterium *Faecalibacterium prausnitzii* through diffusing flavins, which hold significant implications for next-generation probiotics ^7, 20^. Although protein-bound flavins can directly participate in EET without diffusion, soluble shuttling flavins dominate EET processes when their concentration exceeds a certain threshold (e.g., approximately 10 μM in *S. oneidensis*) ^21, 22^. As most cells in biofilms are not in direct contact with electrodes or metal oxides, the diffusion of flavins substantially expands the EET-effective zone in biofilms by transferring electrons over the cellular length ^23^. Notably, synthetic biology approach-based enhancement of flavin secretion has boosted the current output of microbial fuel cells, which highlights the importance of flavin diffusion in biofilms^24^. However, despite its central role, the actual diffusion efficiency of flavins within biofilms remains poorly characterized. A key reason for this knowledge gap is the widespread presumption that diffusion of polar molecules like flavins in biofilms is severely hindered, and their diffusivity is typically considered to be approximately one order lower than that in water ^25, 26^. Based on this assumption, we have not directly quantified flavin diffusivity within this system. Furthermore, the lack of methodological frameworks to evaluate molecular diffusion in biofilms in situ in the presence of potential bias is an additional complication.

In this study, we used interdigitated electrode arrays (IDAs) to evaluate the diffusivity of flavins in electroactive biofilms. IDA-based methods are uniquely suited to extract electron transport properties across biofilms without contributions from intracellular metabolic electron sources. These have been previously used to study the conductivity of outer membrane *c*-type cytochromes and bacterial nanowires in *Shewanella* and *Geobacter* ^27, 28, 29, 30, 31^. However, these approaches have rarely been applied to examine the diffusion of redox molecules in biofilms. Here, we show that contrary to prevailing expectations, the diffusion coefficient of flavins in *S. oneidensis* MR-1 biofilms did not decrease but rather increased up to 11-fold compared with that in bulk solution. Disruption of the *omcA* gene reversed the enhancement, which suggests that the flavin–cytochrome interaction on cell surfaces facilitates flavin diffusion within the biofilm matrix. Furthermore, other redox molecules that shuttle in biofilms, such as methylene blue and safranin, showed scarce enhancement of their diffusion efficiency, which suggests that the biofilm environment specifically enhances flavin diffusion. In summary, this study identifies the previously unrecognized phenomenon that flavin diffusion is assisted by cell-surface *c*-type cytochromes. This could potentially contribute to understanding the physiology, ecology, and energetics of electroactive bacteria in biofilms.

## Results

### Capturing FMN diffusion-mediated electron transfer between interdigitated electrodes

We evaluated flavin diffusion-dependent electron transfer in biofilms by placing a colony biofilm of *S. oneidensis* MR-1 on the IDA with a 10-μm gap between bands (Figure S1). A potential bias ranging from −0.5 to 0.5 V was applied across the electrodes (Figure 1A). The colony was grown on LB agar containing 50 μM flavin mononucleotide (FMN), which is a concentration at which FMN diffusion dominates the electron transfer mechanisms in *S. oneidensis* MR-1 ^21, 32^. We isolated the diffusional contributions of FMN between electrodes by halting metabolic activity to suppress the microbial catalytic current. This was achieved by exposing the colony to erythromycin (4.0 mg mL^−^¹) 1 h before electrochemical measurement, as previously reported ^30, 31^. The *S. oneidensis* MR-1 colony on FMN-containing agar showed a clear current–voltage (I–V) response, and the I/V slope was significantly greater than that in the absence of FMN (Figure 1B). As the I/V slope reflects the conductance of the colony biofilm ^27, 33^, this increase indicates that FMN enhanced the electron transfer between the two interdigitated electrodes separated by the 10-μm gap. Based on a previously reported calculation ^27, 30^, the conductivity (a) of the colony grown on FMN-containing agar was determined to be 9.8 ± 1.4 nS cm^-1^, which was approximately 4.5-fold higher than that without FMN (Figure 1C).

**Figure 1.**
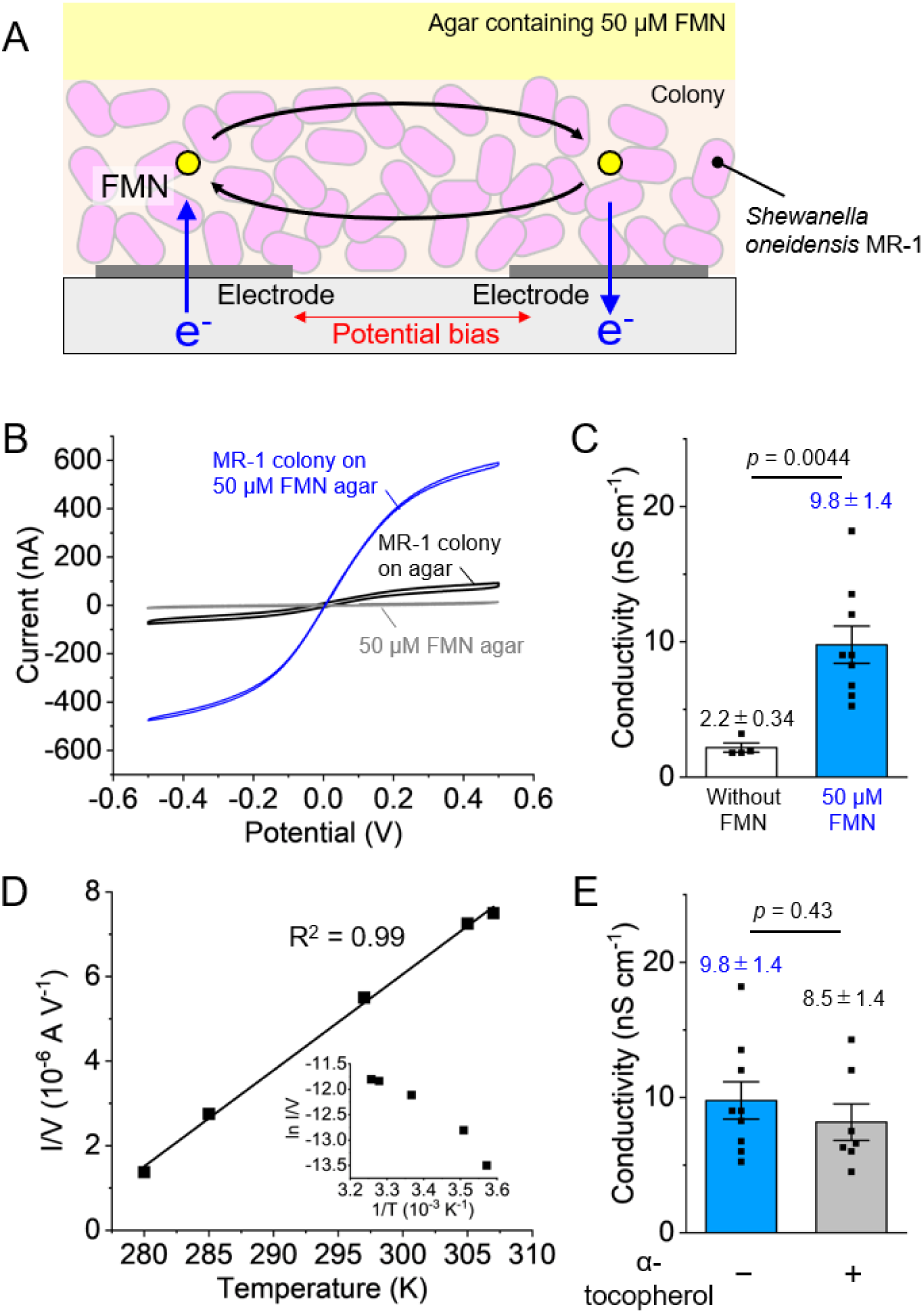
Electron delivery between two electrodes by flavin mononucleotide (FMN) in the colonies of *Shewanella oneidensis* MR-1. (A) A schematic of the experimental setup. MR-1 colonies grown on LB agar containing 50 μM FMN were placed on an interdigitated electrode array (IDA), where a potential bias was applied. (B) Representative current–voltage (I–V) profiles of MR-1 colonies. The colonies were exposed to 4.0 mg mL^−1^ erythromycin before electrochemical measurements to minimize metabolic current generation from MR-1 cells. (C) Effect of 50 μM FMN on electrical conductivity of MR-1 colonies. Error bars represent the standard error of the mean (SEM). Statistical significance was determined based on P-values from unpaired two-tailed Student’s t tests. The data pertaining the MR-1 colony without FMN (black line in B and white bar in C) were obtained from ref. ^30^. (D) Representative plots of I/V vs. temperature for MR-1 colonies cultured on 50 μM FMN-containing agar. They confirmed the temperature-dependent response. I/V was obtained from the slope at −50 to 50 mV of the I–V curve. The square of the correlation coefficient (R^2^) was 0.99, including the point of origin. Inset: Arrehnius-style plot obtained from the identical I–V curve. The same tendency was reproduced in three separate experiments. (E) Effect of α-tocopherol on the I/V slope of MR-1 colonies cultured on 50 μM FMN-containing agar. The error bar represents the S.E.M., and the statistical significance is determined using P-values from unpaired two-tailed Student’s t tests.

We confirmed that the observed current was primarily driven by FMN diffusion by measuring the I–V at various temperatures. If molecular diffusion is the rate-limiting step, according to Fick’s law (equation 1) and Einstein relation regarding Brownian motion (equation 2), the diffusive flux *J* scales proportionally with temperature (*J* ∝ *T*).

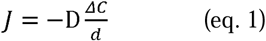

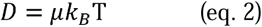

where, D, ΔC, d, μ, and k_B_ indicate the diffusion constant, FMN concentration difference, distance between two electrodes, mobility, and Boltzmann constant, respectively. Thus, I–V profiles were obtained at multiple temperatures under the 50-μM FMN condition (Figure S2). Additionally, a plot of I/V (∝ *J*) against *T* yielded a strong linear correlation (R² = 0.99), which indicates that FMN diffusion is the rate-limiting step for the electron transfer between electrodes in this system (Figure 1D).

Furthermore, the temperature dependence confirmed that cytochrome-based electron-hopping mechanisms did not primarily drive the electron transfer in the colony. A previous study showed that in the absence of flavins, electrons are delivered via cytochromes on the outer membranes of *S. oneidensis* MR-1 under non-diffusion-limited conditions by following an Arrhenius-type behavior (equation 3) ^29^.

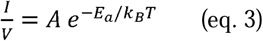

where, A and E_a_ indicate the temperature-independent factor and thermal activation energy, respectively. As Arrhenius plots are temperature-dependent, as shown in equation 3, we plotted *ln(I/V)* against *1/T* (Figure 1D Inset). The fit to the Arrhenius model was weaker than that for the diffusion model. This supports the conclusion that electron transfer between electrodes in the measured system was not mediated by electron-hopping through cytochromes but was rather limited by FMN molecular diffusion.

We further validated the predominance of FMN diffusion by using a radical scavenger. Flavins contribute to EET not only via diffusion but also by binding to outer membrane cytochromes ^30, 31^. In these cases, flavin is present in a one-electron reduced semiquinone state, which is quenched by the radical scavenger α-tocopherol ^21, 31^. Hence, we added α-tocopherol to an *S. oneidensis* MR-1 colony at a sufficient concentration (5.0 mM) to quench semiquinone ^31^; however, the I/V slope showed only a mild decrease, which corresponded to only a 13% decrease in conductivity (Figures 1E and S3). This finding indicates that the majority of electron transfer observed in this system with 50 μM FMN, was mediated by the diffusing FMN rather than the FMN–cytochrome complex.

### Increase in FMN diffusion coefficient in *S. oneidensis* MR-1 biofilms

We calculated the diffusion coefficient of FMN in the colony biofilm according to the flowchart shown in Figure 2A. Specifically, the apparent diffusion coefficient in biofilm (D_biofilm_) was calculated using the Nernst–Einstein equation (equation 4), which links the conductivity a with the diffusion coefficient.

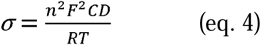

where, n, F, C, and R indicate the number of electrons involved in the FMN redox reaction (= 2), Faraday constant, FMN concentration, and gas constant. Although the colony was cultured on agar containing 50 μM FMN, the actual FMN concentration within the colony may differ. Therefore, we estimated the colony volume through microscopic observation and quantified the FMN quantity by measuring the fluorescence intensity after dispersing it in PBS (Figures 2A, S4, S5, and S6). The FMN concentration in the colony was 33 ± 4.0 μM (Figure 2B). Using this concentration, the D_biofilm_ was calculated to be 1.9 ± 0.27 × 10^-9^ m^2^s^-1^ (Figures 2C-E). The FMN diffusion coefficient in agar was 4.2 ± 1.6 × 10^-12^ m^2^s^-1^, which was significantly lower than the D_biofilm_ (Figure S7). This indicates that the FMN diffusion coefficient is increased in the colonies. We further validated this finding by quantifying the diffusion coefficient of FMN in the LB medium solution based on the I–V measurement and using chronoamperometry, which is a well-established approach to estimate diffusion coefficients in solution ^34^. Chronoamperometry was performed using a popular three-electrode system for measurement ^35^. The diffusion coefficients were determined to be 1.1 ± 0.05 ×10^-11^ m^2^s^-1^ and 0.17 ± 0.01 ×10^-9^ m^2^s^-1^ (Figures 2E, S6, and S7) based on the I–V measurement and chronoamperometry, respectively. The latter value was closer to the reported FMN diffusion coefficient in aqueous solution (0.38 ×10^-9^ m^2^s^-1^) ^36^, which is probably because chronoamperometry is suited for a direct determination of diffusion coefficients in solution through the isolation of the diffusion-limited current predicted by the Cottrell equation, whereas the IDA-based system favors probing lateral shutting behavior within confined colonies ^30^. These results show that the FMN diffusion coefficient in the colony biofilm is at least 11-fold higher than that in the solution.

**Figure 2.**
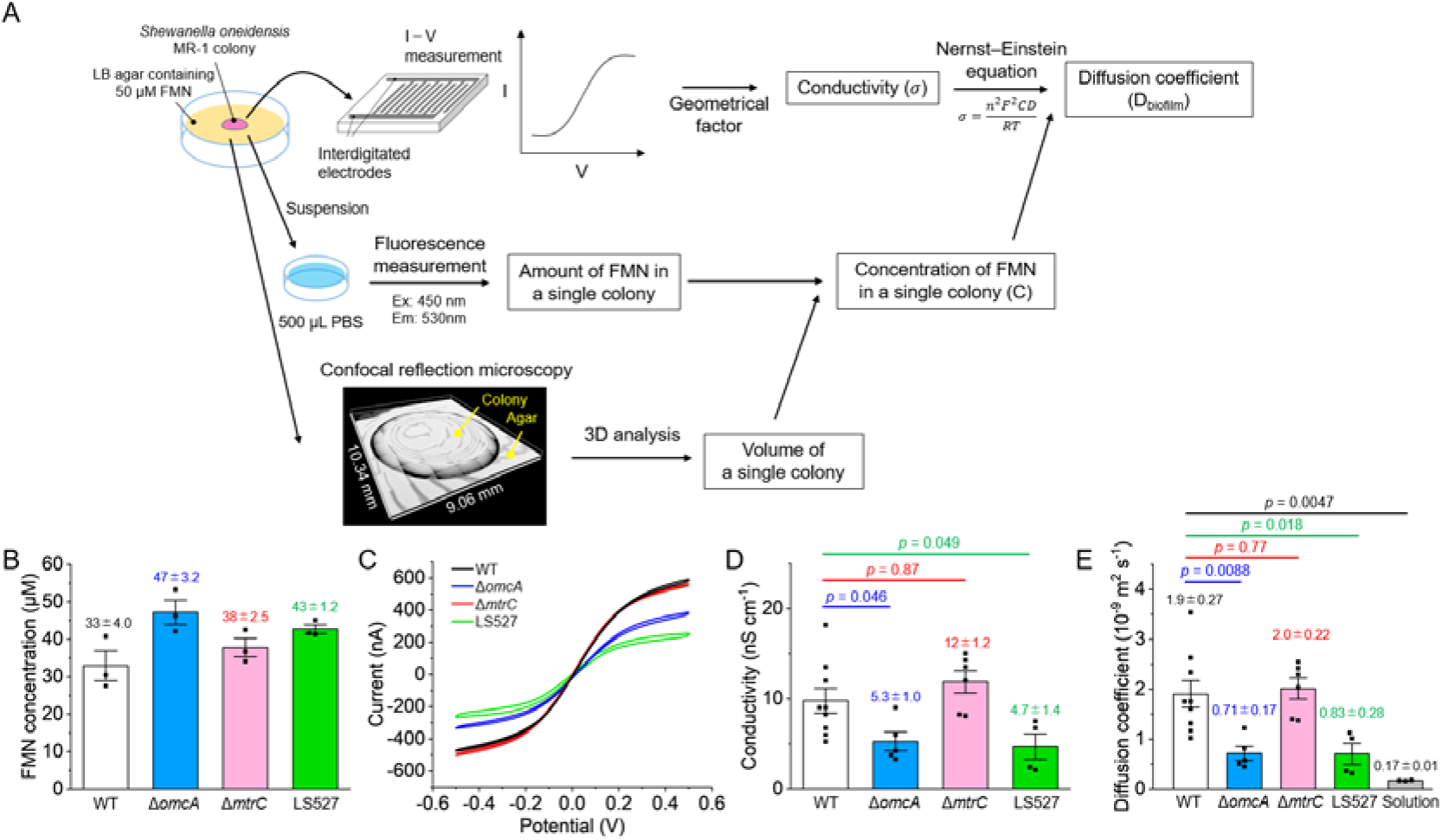
Promotion of electron delivery in *S. oneidensis* MR-1 colonies by shuttling flavins, and its relationship with cytochromes. (A) Schematic of experimental flowchart to quantify the FMN diffusion coefficient in MR-1 colonies (D_biofilm_). (B) Concentration of FMN in MR-1 gene deletion mutant colonies cultured on 50 μM FMN agar. (C) Representative I–V profiles of MR-1 gene deletion mutant colonies cultured on 50 μM FMN agar. Data of wild type (WT, black line) were identical to that shown in Figure 1B. (D) Conductivity of MR-1 colonies cultured on 50 μM FMN agar. (E) FMN diffusion constants. Each plot was calculated from each conductivity of the colony using the averaged FMN concentration in colonies. Diffusion constants in aqueous solution obtained from chronoamperometric measurements. The error bar represents the S.E.M., and statistical significance was determined based on P-values obtained using unpaired two-tailed Student’s t tests.

### A *c*-type cytochrome is involved in the increase of FMN diffusion coefficient in *S. oneidensis* MR-1 biofilms

We identified the cause of increase in the diffusion coefficient in the colonies by measuring I–V using gene deletion mutants. As outer membrane *c*-type cytochromes interact with the flavins in the biofilms ^30, 31^, we hypothesized that these interactions underlie the observed enhancement. We cultured colonies of Δ*omcA* and Δ*mtrC* mutants that lacked the respective outer membrane *c*-type cytochrome on agar containing 50 μM FMN, and I–V measurements were performed under the same conditions (Figure 2B). The I/V slope showed a pronounced decrease in the Δ*omcA* colony, whereas the change was negligible in the Δ*mtrC* colony compared with that of the wild type (WT). The same trend was observed when converted to conductivity: only Δ*omcA* showed a significant decrease (WT: 9.8 ± 1.4 nS cm^-1^, Δ*omcA*: 5.3 ± 1.0 nS cm^-1^, and Δ*mtrC*: 12 ± 1.2 nS cm^-1^) (Figure 2C). Additionally, the diffusion coefficient was significantly decreased only in Δ*omcA* (WT: 1.9 ± 0.27 × 10^-9^ m^2^s^-1^, Δ*omcA*: 0.71 ± 0.17 × 10^-9^ m^2^s^-1^, and Δ*mtrC*: 2.0 ± 0.22 × 10^-9^ m^2^s^-1^) (Figure 2D). These results strongly suggest that OmcA facilitates FMN diffusion in the *S. oneidensis* MR-1 biofilm.

Additionally, a strain lacking multiple cytochromes, including *mtrCAB* (SO_1776–SO_1778), *mtrDEF* (SO_1780–SO_1782), and *omcA* (SO_1779), termed LS527 ^37^, exhibited a significantly lower diffusion coefficient than that of the WT (0.83 ± 0.28 × 10^-9^ m^2^s^-1^). However, both the colony volume and total cell number within the colonies of these mutants were comparable with those of the WT (Figures S6 and S9), which suggests that the observed differences in diffusion cannot be attributed to differences in cellular density in the biofilm.

### Diffusion coefficient increases specifically for FMN

We determined whether the observed increase in diffusion coefficient was specific to FMN or a general property of redox molecules by extending this analysis to other representative electron shuttles that are compatible with *S. oneidensis* MR-1 such as methylene blue (MB) and safranin (SF)^38^. Colonies were cultured on agar supplemented with 50 μM MB or SF, and I–V measurements were performed. Similar to that observed for FMN, both MB and SF showed significant increase in the I/V slope, which indicates that these redox molecules also facilitate long-distance electron transfer in MR-1 colonies (Figure 3A). The calculated conductivities were 39 ± 3.6 nS cm^-1^ for MB and 6.2 ± 0.51 nS cm^-1^ for SF (Figure 3B), which is indicative of their high efficiency in promoting long-range electron transport.

**Figure 3.**
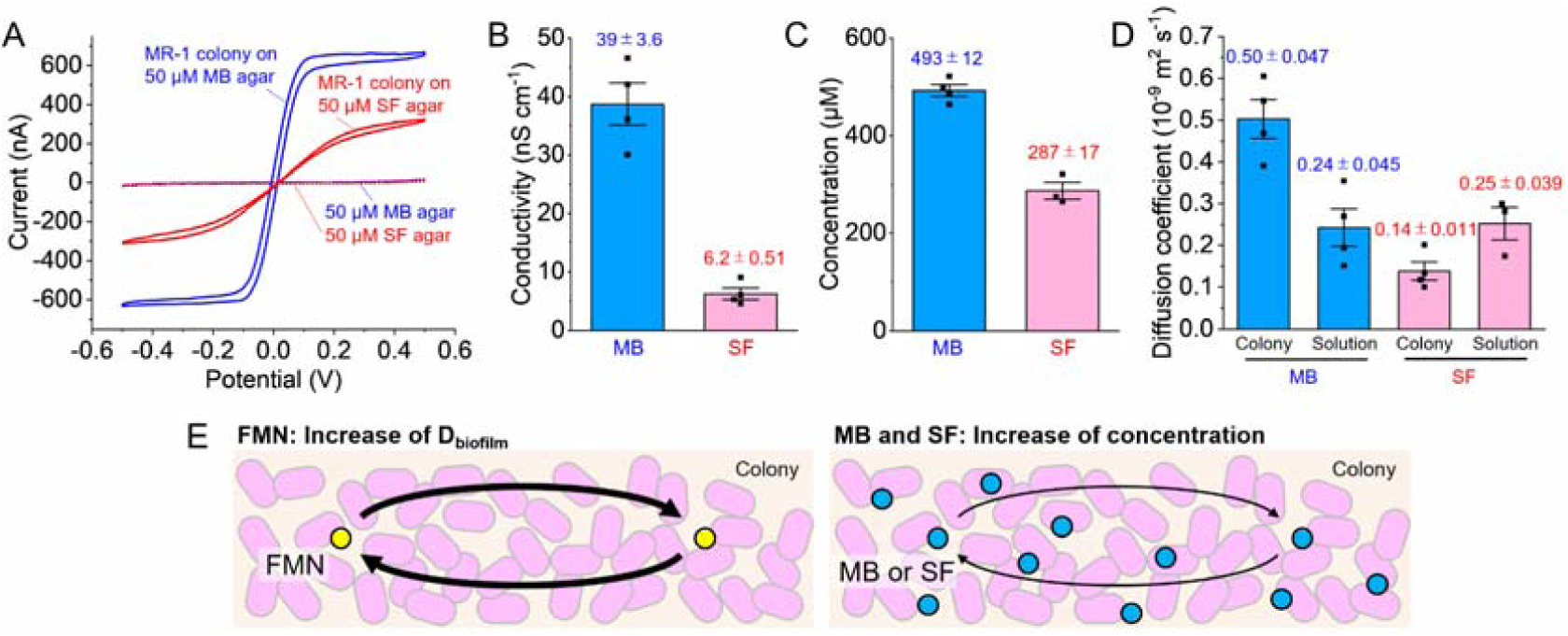
Electron delivery in *S. oneidensis* MR-1 colonies through methylene blue (MB) and safranin (SF) shuttling. (A) Representative I–V profiles of MR-1 colonies grown on LB agar containing 50 μM MB or SF. (B) Conductivity of MR-1 colonies cultured on 50 μM MB or SF agar. (C) Concentration of MB or SF in MR-1 colonies. (D) Diffusion coefficients of MB and SF in MR-1 colonies or medium solution. Each plot was calculated from each conductivity of the colony using the averaged MB or SF concentration. The error bar represents the S.E.M., and statistical significance was determined using P-values from unpaired two-tailed Student’s t tests. (E) We constructed two models for the promotion of electron delivery in MR-1 colonies: FMN enhances electron transfer by increase in D_biofilm,_ and MB and SF enhance electron transfer by increased MB or SF accumulation inside the colonies.

Notably, MB and SF concentrations within the colonies were significantly higher than the initial 50 μM in the agar and reached 493 ± 12 μM and 287 ± 17 μM, respectively (Figure 3C). This phenomenon was similar to the accumulation of pyocyanin as a mechanism to facilitate long-range electron transport in *Pseudomonas aeruginosa* colonies ^39^. However, the MB diffusion coefficient in biofilm did not show more than a 2.0-fold increase, and SF diffusion coefficient in biofilms decreased (Figure 3D). This shows that the enhancement of electron transfer for MB and SF is primarily driven by their accumulation within the colony rather than by changes in diffusivity. Furthermore, the I–V profiles were scarcely affected by gene deletion for cytochromes (Figure S10). Thus, we inferred that the OmcA-assisted one-order enhancement of D_biofilm_ observed for FMN was not a general feature of redox molecules but rather a unique property of FMN in MR-1 biofilms. These findings show that distinct redox molecules promote long-distance electron transfer through different mechanisms within *S. oneidensis* MR-1 biofilms. FMN does so by increasing diffusivity, whereas MB and SF function via concentration-dependent enhancement (Figure 3E).

## Discussion

Electroactive bacteria that utilize extracellular solid materials as terminal electron acceptors face unique ecological challenges compared with those faced by bacteria that utilize soluble acceptors. Generally, reduction reactions occur only at heterogeneous two-dimensional surfaces; hence, they are inherently limited in terms of spatial accessibility to terminal electron acceptors. Although the formation of three-dimensional biofilms offers numerous advantages such as protection against environmental fluctuations and toxic compounds, the mechanism underlying the efficient access of solid electron acceptors by electroactive bacteria within these spatially structured communities remains unclear. *Geobacter* is a model electroactive bacterium that addresses this challenge using electrically conductive biofilms that enable EET participation from cells located far from the electrode. Although *Geobacter* possesses the highest known conductivity, its EET activity significantly decreases farther than approximately 10 µm from the electrode surface and becomes negligible at distances more than approximately 45 µm ^40, 41^. Typical bacterial conductivities are several orders lower than that of *Geobacter* ^42, 43^; hence, conduction-based EET may be reasonably assumed to be insufficient to support metabolic activity in most biofilms. In contrast, molecular diffusion is often recognized as kinetically less favorable than conduction, and biofilm matrices are generally thought to impede diffusion ^25, 26^. Thus, the mechanism underlying the ability of the electroactive bacteria that are not in direct contact with solid acceptors to sustain metabolic activity via EET remains unclear. The present study answers this long-standing ecological puzzle by showing that diffusion of flavins is notably enhanced in the biofilm matrix. This enhancement contradicts conventional assumptions as it provides a solution to overcome the spatial limitations faced by electroactive bacteria by enabling access to terminal electron acceptors. Moreover, this enhancement appears to be specific to flavins, as evidenced by the lack of significant enhancement of diffusivity of artificial redox molecules of similar molecular weight such as MB and SF. This specificity reinforces the notion that *S. oneidensis* has adapted to optimize the efficiency of its own secreted flavins.

Furthermore, this flavin diffusion enhancement highlights the importance of considering the physicochemical behavior of redox molecules within biofilm environments when designing EET-based bioelectronic devices such as microbial fuel cells and microbial electrosynthesis. The selection of redox mediators for such applications has largely been based on measurements involving planktonic cells or cells adsorbed on electrodes ^44^. The current study introduces biofilm-localized diffusivity as a new evaluation axis for redox mediators. The development of flavin derivatives could be a powerful approach for identifying high-performance redox molecules for EET-based applications.

Additionally, we have elucidated a mechanistic basis for the enhancement of FMN diffusion by identifying that OmcA, which is a cell-surface flavin-binding cytochrome, facilitates flavin diffusion within the biofilm. Furthermore, the increase in D_biofilm_ was observed under metabolically inactive conditions in our experimental setup, implying that the enhancement is not driven by redox cycling or active transport. The FMN concentration in this study was 50 µM, which is at least 1–2 orders lower than the concentration that enables electron hopping as observed for quinones ^45^, which is also confirmed by the temperature dependence (Figure 1D). Therefore, we propose a model in which flavin diffusion is promoted via surface diffusion along the cell envelope. FMN binds to OmcA with moderate affinity (with reported dissociation constants ranging from approximately 10 to several hundred μM ^21, 22, 46^) at the same range as the FMN concentration used in our experiments. This suggests a dynamic equilibrium between membrane-bound and free flavins within the biofilm. Diffusion is a phenomenon where a ligand that is loosely associated with membrane proteins or surfaces migrates along the interface. Diffusion has been reported in some biological systems ^47, 48, 49^; however, it has rarely been considered in the context of microbial EET. Although current methodologies do not yet permit the quantitative modeling of the impact of surface diffusion on electron flux, such spatial confinement could conceivably enable electron delivery to electrodes with better efficiency than that of free three-dimensional diffusion. Future studies that use single-molecule tracking techniques within biofilms are needed to enable the direct visualization and quantification of this proposed surface diffusion.

Notably, the FMN diffusion coefficient remained higher than that in aqueous solution even in the cytochrome-deficient *S. oneidensis* strain LS527, which lacked multiple outer membrane cytochromes, including OmcA. This suggests that additional unidentified factors may contribute to the enhanced flavin diffusion observed in MR-1 biofilms. Systematic approaches such as transposon mutagenesis or proteomic screening may help identify novel components that modulate diffusion dynamics within microbial communities.

In conclusion, we have successfully quantified the apparent diffusion coefficient of redox molecules within bacterial colonies by combining colony-based electrochemistry, conductivity measurements using IDAs, and calculations based on the Nernst–Einstein equation. Our results show that *S. oneidensis* MR-1 possesses mechanisms that enhance flavin-mediated electron shuttling in biofilms through OmcA–flavin interaction. This finding challenges previous assumptions regarding the diffusional limitations of biofilms and establishes a new framework for understanding and engineering electroactive biofilms in natural and applied settings. The elucidation of the molecular and biophysical principles that govern flavin behavior in biofilms provides foundational knowledge for the rational design of next-generation bioelectronic devices and offers insight into microbial bioenergetics in biofilm environments.

## Methods

### Bacterial strain and colony growth conditions

An *S. oneidensis* MR-1 colony cultured on Luria–Bertani (LB) agar [NaCl (5.0 g L^−1^), yeast extract (5.0 g L^−1^), tryptone (10 g L^−1^), and Bacto agar (15 g L^−1^)] was picked and transferred into 4.0 mL of LB medium and aerobically cultured overnight at 30 °C and with shaking at 190 rpm. Then, the culture was centrifuged at 6,000 × *g* for 5 min. The obtained cell pellet was resuspended in LB at an optical density at 600 nm of 0.25. A 10-μL cell suspension was dropped on LB agar containing 50 μM FMN and cultured for 24 h at 30 °C to form a colony. Before performing electrochemical measurement, the colony was treated with erythromycin (4 mg mL^-1^) and left for 1 h to suppress microbial catalytic current by halting metabolic activity, as previously reported ^30, 31^. The mutant strains Δ*mtrC,* Δ*omcA* (reported in ref ^21^), and LS527 (reported in ref ^37^) were used.

### Current–voltage (I–V) measurement of bacterial colonies with interdigitated electrode arrays (IDAs)

We used the IDAs reported previously ^30, 31^. The IDAs comprised 704 parallel indium tin-doped oxide bands (352 pairs) that were 10 mm long, 10 μm wide, separated by 10-μm gaps, and patterned onto a glass substrate (Geomatec Co. Japan). Each adjacent band was electrically connected to the opposite ends of the array to form two interdigitated electrodes. A colony on agar was placed on the IDA to directly face the interdigitated electrodes with the colony. Potential bias was applied between the interdigitated electrodes from −0.5 to 0.5 V at room temperature (25 °C) aerobically at a scan rate of 1.0 mV s^−1^ to minimize the capacitive charging current using automatic polarization systems (VMP3, BioLogic Co., France). To evaluate the temperature dependence of the I–V curves, the scan rate was increased to 10 mV s^−1^ to minimize time-dependent changes in the sample during prolonged measurements. The potential bias was applied for three cycles, and the third cycle was shown.

### Three-dimensional imaging and quantification of bacterial colony volumes using confocal reflection microscopy

Bacterial colonies were visualized using an LSM880 confocal laser scanning microscope equipped with a Plan-Apochromat dry objective lens at 10× magnification and a 0.45-numerical aperture (Carl Zeiss, Jena, Germany). The colonies were irradiated with a 633-nm laser, and reflections in the 625–646-nm range were used to construct confocal reflection microscopic images ^50^. The images were obtained through tile scanning at a z-scanning resolution of 10.00 μm.

Colony volume was quantified using ImageJ and R by following a multi-step topographical reconstruction workflow (Figure S5). First, three-dimensional colony topography was generated from z-stack fluorescence images using the TopoJ plugin in ImageJ. Next, the colony region was manually selected and masked to produce an image representing the agar substrate beneath the colony. Next, an interpolated surface of the underlying agar was reconstructed using the Akima interpolation method implemented in R ^51^. Finally, colony-specific height values were obtained by subtracting the interpolated agar surface from the original topographical dataset, and the total colony volume was calculated from the obtained height map.

### Calculation of diffusion coefficients in bacterial colonies

As shown in Figure 2A, diffusion constants of redox molecules in bacterial colonies were calculated from the Nernst–Einstein equation (equation 4) using electrical conductivity (σ) and concentration of redox molecules in the colonies.

Colony conductivity was determined based on a previously reported methodology ^30^ as follows:

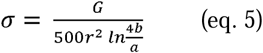

where, *G* and r indicate the conductance and radius of the colony. a and b were considered 5 μm and 10 μm, respectively, as defined based on the geometry of the IDA, as reported ^52^. As the I–V slope close to 0 V represents the conductance ^27, 33^, the conductance was obtained from the slope at −50 mV to 50 mV by following Ohm’s Law.

The concentration of redox molecules within the colonies was estimated by combining microscopic measurements of colony volume with fluorescence-based quantification of redox molecules. To quantify redox molecules in colonies, colonies were suspended in 500 μL PBS and subsequently applied to a microplate reader (BioTek SYNERGY H1 microplate reader, Winooski, VT). The following excitation and emission wavelengths were used for fluorescence measurements: flavin, 450 nm/530 nm; MB, 650 nm/700 nm; and SF, 500 nm/590 nm. By dividing the amount of redox molecules by the colony volume, the concentrations of redox molecules within the colonies were provided.

### Diffusion constant measurement of FMN without bacterial colonies

Diffusion coefficients of FMN in LB solution were determined using two approaches: 1) calculation of diffusion coefficients using the I–V curves and 2) calculation of diffusion coefficients based on chronoamperometric measurements. Calculating the diffusion coefficients from the I–V curves involves the assumption that FMN is confined near the electrodes ^30^; hence, applying this method to freely diffusing molecules in solution potentially underestimates the diffusion coefficients. Therefore, to accurately determine the diffusion coefficients of FMN in bulk solution, we used a well-established chronoamperometry-based method that assumes free diffusion in solution ^34^. Chronoamperometry was performed using a three-electrode system, in which an ITO working electrode (surface area: 3.1 cm²) was used, as previously reported ^35^. Each redox molecule (FMN, MB, and SF) was added into LB medium at a final concentration of 50 μM under anaerobic conditions. After applying a constant potential that was sufficiently positive compared with the redox potential of the redox molecules (FMN, +0.1 V; MB, +0.25 V; and SF, –0.15 V vs. SHE) for 30 s, the potential was stepped to a sufficiently negative value (FMN, –0.4 V; MB, –0.2 V; and SF, –0.6 V vs. SHE). The resulting reduction current was recorded at 50 ms intervals, and the diffusion-limited current between 1.0 and 2.0 s was used to calculate the diffusion coefficient based on the Cottrell equation as described:

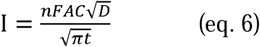

## Acknowledgements

We thank Prof. Dr. Liang Shi for kindly providing the LS527 mutant. This work was financially supported by JST ACT-X (JPMJAX211C); JST FOREST (JPMJFR231H); JST GteX (JPMJGX23B2); and JSPS KAKENHI (20K15428 and 23H05471).

## Contributions

Y. T. designed and supervised the project and wrote the manuscript. Y. T. and Y. M. conducted the experiments and performed the data analyses. All authors contributed to reviewing and editing the manuscript with constructive discussions.

## Data availability

All data supporting the findings of this study are available within the main text, supplementary information, and source datasets. The source datasets are available in Figshare (https://doi.org/10.6084/m9.figshare.30782939).

