## Supplementary file for "Challenging the Diffusion Barrier Paradigm: Biofilms Promote Flavin-mediated Electron Shuttling in *Shewanella oneidensis*"

**in *Shewanella oneidensis***

Yoshihide Tokunou^1, 2^*, Yu Manabe^3^, Nozomu Obana^4, 5, 6^, Masanori Toyofuku^1, 4, 5^, and Nobuhiko Nomura^1, 4, 5^

^1^ Department of Life and Environmental Sciences, University of Tsukuba, Tsukuba, Ibaraki 305-8577, Japan

^2^ Research Center for Macromolecules and Biomaterials, National Institute for Material Science, 1-2-1 Sengen, Tsukuba, Ibaraki, 305-0047, Japan

^3^ Degree Programs in Life and Earth Sciences, University of Tsukuba, 1-1-1 Tennodai, Ibaraki 305-8577, Japan

^4^ Microbiology Research Center for Sustainability, University of Tsukuba, Tsukuba, Ibaraki 305-8572, Japan

^5^ Tsukuba Institute for Advanced Research, University of Tsukuba, Tsukuba, Ibaraki 305-8572, Japan

^6^ Department of Medicine, Transborder Medical Research Center, University of Tsukuba, Tsukuba, Ibaraki 305-8577, Japan

*Corresponding Author: Y. T.

Contents:

Supplementary Figures 1-10

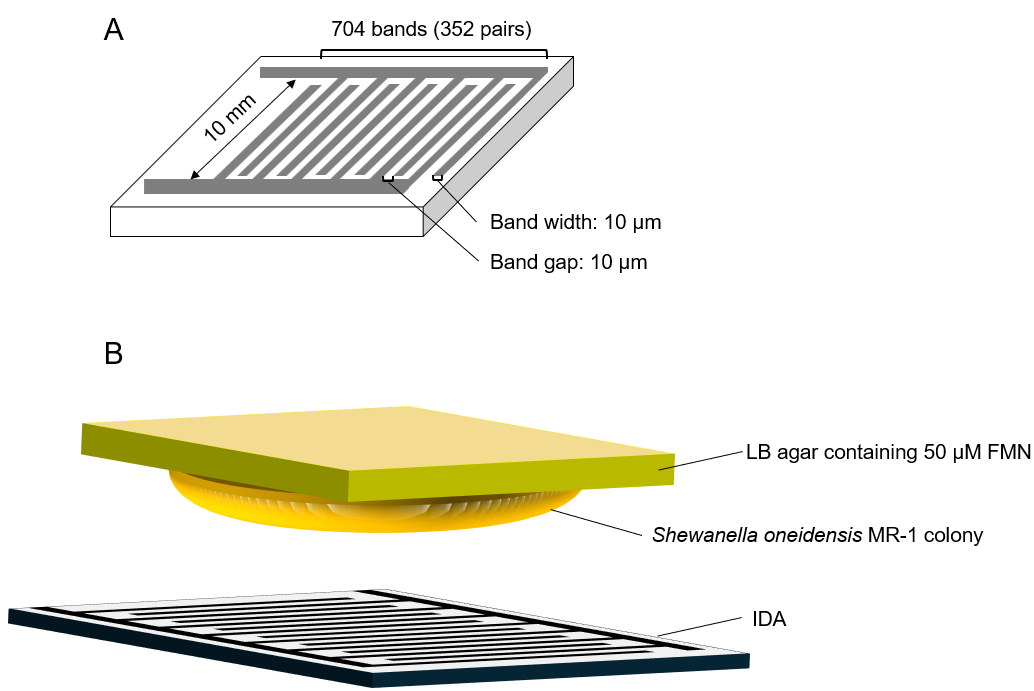

**Figure S1. Schematic illustrations of the detailed experimental setup.** (A) An illustration of an interdigitated electrode array (IDA). Interdigitated electrodes (10 mm long and 10 μm wide) are patterned with 10 μm wide gaps. (B) An illustration of electrochemical measurement using *Shewanella oneidensis* MR-1 colony biofilm and IDA.

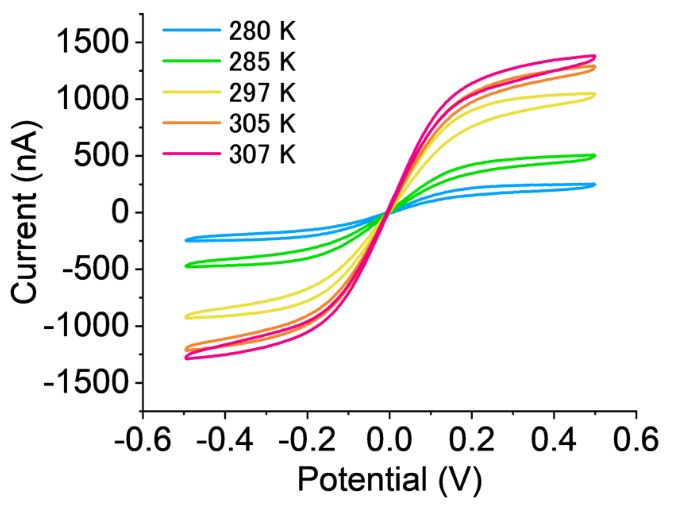

**Figure S2. Temperature-dependence of current–voltage (I–V) profiles of *Shewanella oneidensis* MR-1 colonies on 50 μM FMN agar.**

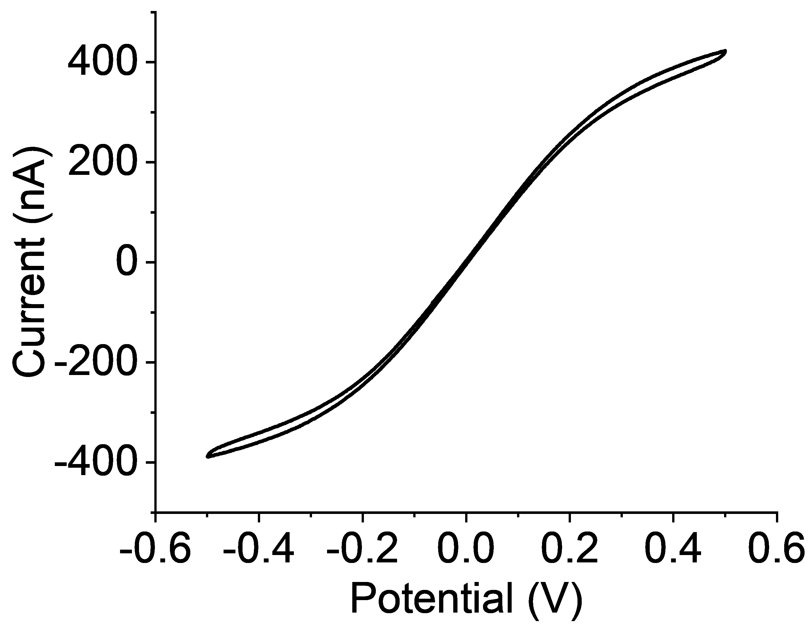

**Figure S3. A representative I-V curve of *Shewanella oneidensis* MR-1 colonies on 50 μM FMN agar in the presence of 5.0 mM α-tocopherol.**

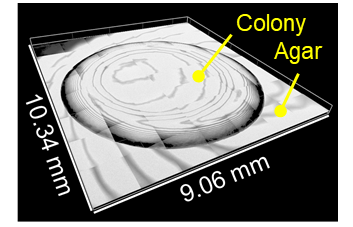

**Figure S4. A representative three-dimensional image of *Shewanella oneidensis* MR-1 colony observed with confocal reflection microscopy.**

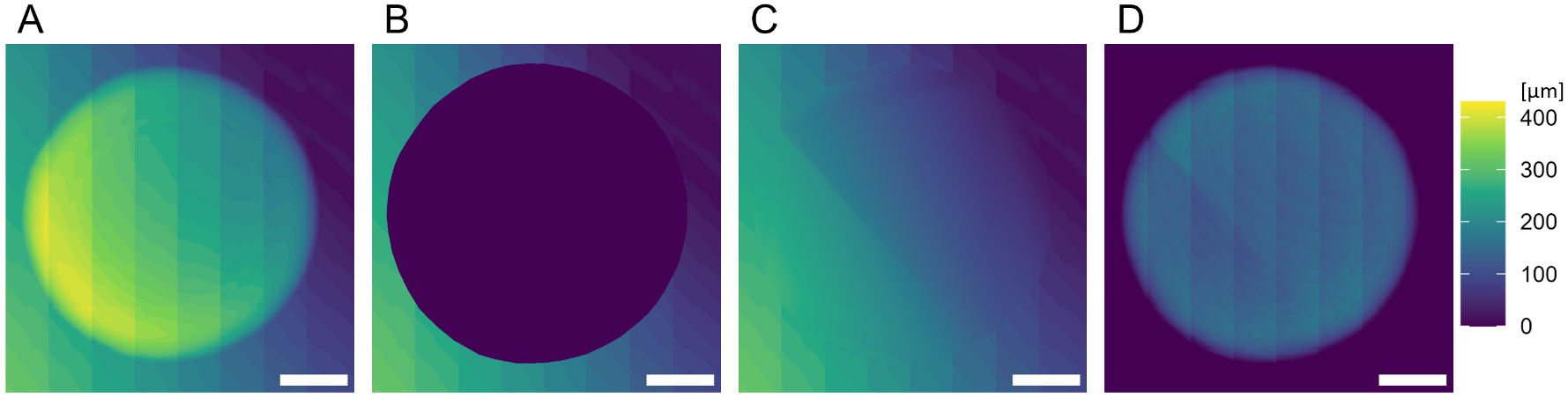

**Figure S5. Representative hight maps of *Shewanella oneidensis* MR-1 colonies for calculation of the colony volume.** (A) The topography generated from the original *Shewanella oneidensis* MR-1 colony image. (B) The topography in which the colony region was manually masked. (C) The interpolated image of the agar beneath the colony. (D) The colony-specific topography generated by subtracting the interpolated image from the original topography. Scale bars indicate 2 mm.

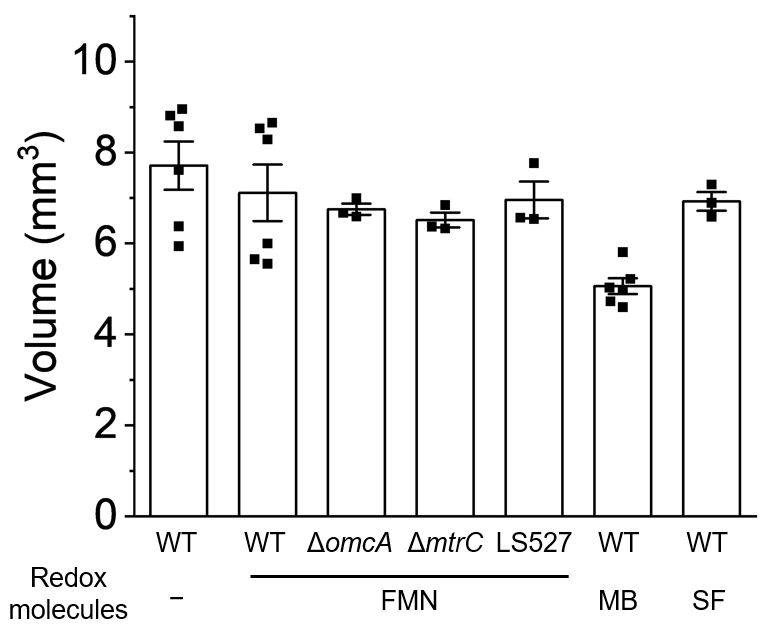

**Figure S6. Volume of *Shewanella oneidensis* MR-1 colonies.** The error bar represents the standard error of the mean (S.E.M.).

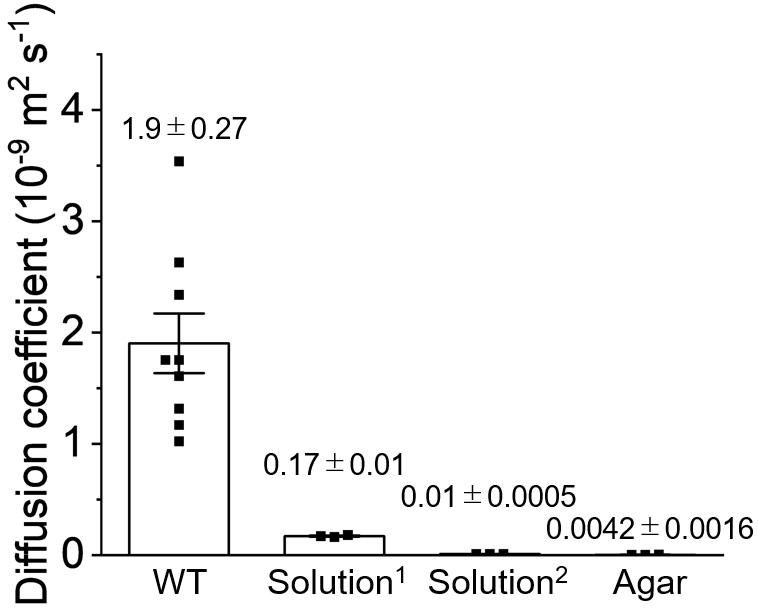

**Figure S7. Diffusion coefficients of FMN in *Shewanella oneidensis* MR-1 colonies, LB solution, and LB agar.** The data for WT and Solution^1^ are identical to those in Figure 2E. The data for Solution^1^ and Solution^2^ represent FMN diffusion constants calculated from chronoamperometric data and I–V curves, respectively. The error bar represents the S.E.M..

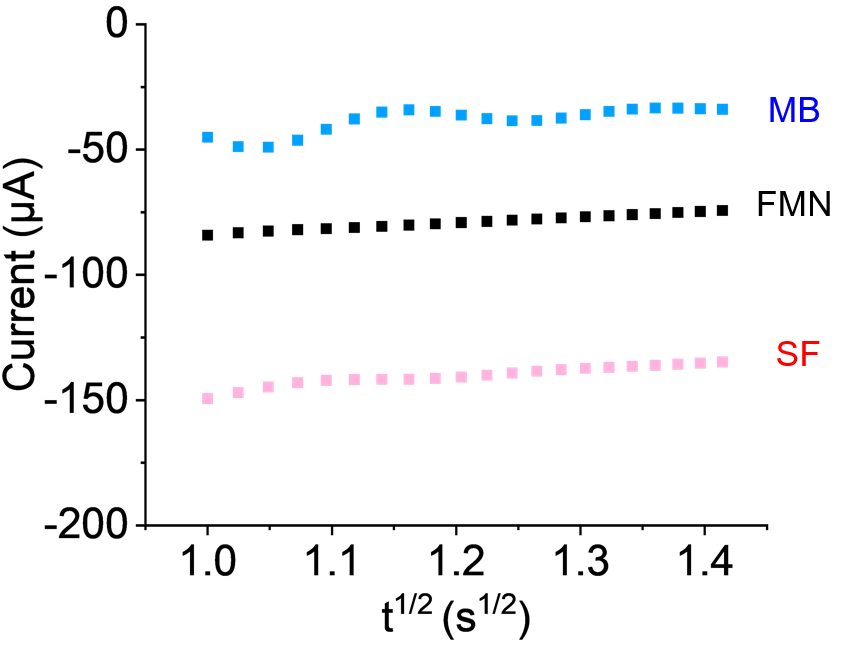

**Figure S8. Representative chronoamperometry data of 50 μM FMN, methylene blue (MB), and safranin (SF) for the determination of diffusion coefficients in solution.** Each redox molecule (FMN, MB, and SF) was added to a final concentration of 50 μM in an anaerobic three-electrode system, and the currents were recorded under constant potential application (FMN, –0.4 V; MB, –0.2 V; and SF, –0.6 V vs. SHE). Before applying a constant potential, enough positive potential compared to the redox potential of each redox molecule was applied (FMN, +0.1 V; MB, +0.25 V; and SF, –0.15 V vs. SHE) for 30 s. The resulting reduction current was recorded at 50 ms intervals, and the diffusion-limited current between 1.0 and 2.0 s was used to calculate the diffusion coefficient based on the Cottrell equation.

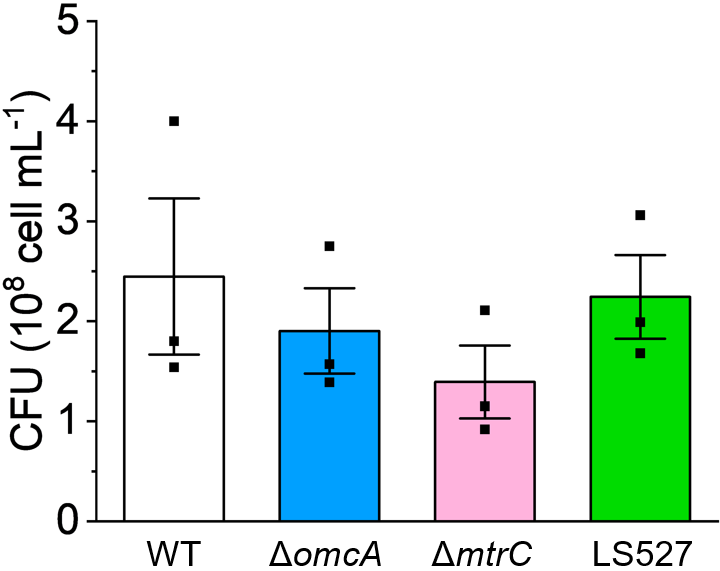

**Figure S9. CFU (colony-forming unit) of a single *Shewanella oneidensis* MR-1 colony on 50 μM FMN agar.** The error bar represents the S.E.M..

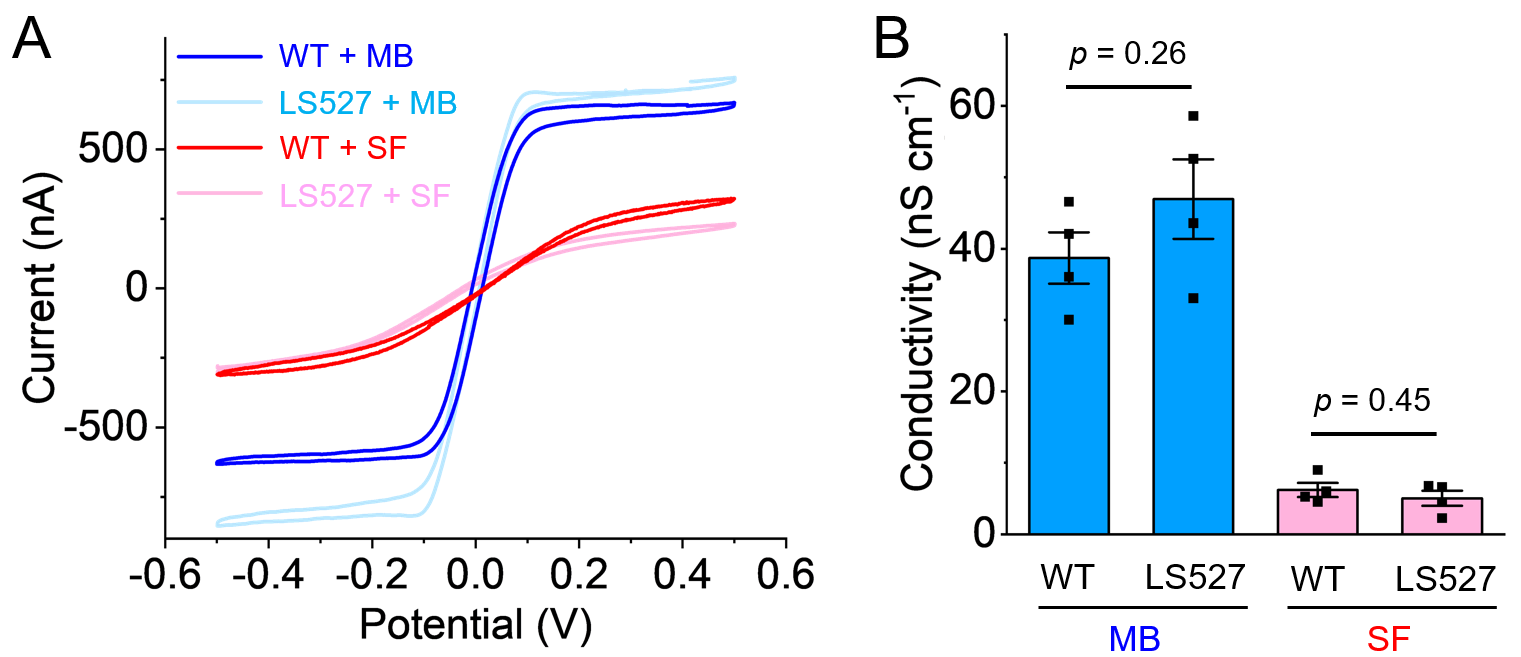

**Figure S10. The impact of methylene blue (MB) and safranin (SF) on LS527 colonies.** (A) Representative I–V profiles of *S. oneidensis* MR-1 WT and LS527 colonies in the presence of 50 μM MB or SF. (B) The electrical conductivity of WT and LS527 colonies in the presence of 50 μM MB or SF. The error bar represents the S.E.M.. Statistical significance was determined based on P-values from unpaired two-tailed Student’s t tests.
